# The Regulation of MAGI2 and its Diagnostic Application in Podocytopathies

**DOI:** 10.1101/2024.12.01.626009

**Authors:** Florian Siegerist, Eleonora Hay, Elke Hammer, Anna Iervolino, Claudia Weber, Juan Saydou Dikou, Eleni Stamellou, Linus Butt, Thomas Benzing, Thorsten Wiech, Paul T. Brinkötter, Uwe Zimmerman, Giovambattista Capasso, Christos Chatziantoniou, Christos Chadjichristos, Tobias B. Huber, Uwe Völker, Maximilian Schindler, Nicole Endlich

## Abstract

Podocyte dysfunction is central to various glomerular diseases, necessitating reliable biomarkers for early detection and diagnosis. This study investigates the regulatory mechanisms of membrane-associated guanylate kinase inverted 2 (MAGI2) and its potential as a biomarker for podocytopathies. The expression of the gene coding for the scaffolding protein MAGI2 was examined across four species and demonstrated to be conserved within the podocyte filtration slit. *In vitro* and *in vivo* studies using isolated glomeruli and mammalian animal models of glomerular disease, including DOCA-salt hypertension, nephrotoxic serum nephritis, and puromycin aminonucleoside nephropathy, demonstrated significant downregulation of MAGI2 in injured podocytes. This downregulation was also conserved in a zebrafish model of focal and segmental glomerulosclerosis (FSGS), and the podocyte-specific MAGI2 ortholog Magi2a was reduced post podocyte injury. CRISPR/Cas9-generated zebrafish mutants for *magi2a* exhibited marked glomerular filtration barrier defects and downregulation of nephrin, underscoring MAGI2’s critical role in podocyte function.

Human biopsy analyses revealed differential MAGI2 expression: it was increased in minimal change disease (MCD) patients but significantly decreased in primary, but not secondary FSGS cases. As MAGI2 localization did not change in disease states it is an alternative marker for super-resolution microscopy-based morphometry of the filtration slit, correlating with nephrin-based measurements.

These findings highlight the potential of MAGI2 as a sensitive biomarker for podocyte injury and its diagnostic utility in differentiating between primary FSGS and MCD.

## Introduction

The glomerular filtration barrier (GFB) is essential for permselective blood filtration and is composed of three layers: endothelial cells, the glomerular basement membrane (GBM), and podocytes. Neighboring interdigitating podocyte foot processes form highly specialized cell-cell contacts that are described ultrastructurally as the slit diaphragm [1]. The slit diaphragm is composed of a multiprotein complex that contains features of adherens-as well as tight junctions which components link the large extracellular domains of nephrin to the actin cytoskeleton [1,2]. Podocyte injury initiates foot process effacement, a process that involves the loss of the highly complex three-dimensional structure of the foot processes, and the impairment of the function of the filtration barrier, characterized by high-molecular-weight proteinuria. Glomerular diseases in which high-molecular-weight proteinuria is attributed to the damage or dysfunction of the podocytes are grouped under the name of podocytopathies [3]. Podocytopathies have multiple reasons (genetic, autoimmune, or maladaptive) and include histological patterns like minimal change disease (MCD) and focal segmental glomerulosclerosis (FSGS).

Previously it has been shown that the podocyte cell-cell contacts contain several tight junction or tight junction-associated proteins like TJP-1, JAM-A, and CLDN5 [2,4,5]. These proteins are upregulated during podocyte foot process effacement, with some being present in the healthy slit diaphragm (e.g. TJP1 [2]) and others being recruited to the cell-cell contacts of effaced podocyte foot processes (CLDN5 [5]). One tight junction-associated protein that has been shown to be localized within the slit diaphragm complex is MAGI2 [4]. The scaffolding protein MAGI2 is a membrane-associated guanylate cyclase that has been shown to be important for kidney barrier function since knockout (KO) mice die early from end-stage kidney failure[6]. In patients, recessive mutations in the MAGI2 gene lead to a steroid-resistant nephrotic syndrome with a focal and segmental sclerosing (FSGS) histologic phenotype indicating primary podocyte injury [7]. Shirata and colleagues generated podocyte-specific MAGI2 KO mice demonstrating that they exhibit glomerulosclerosis and kidney damage similar to FSGS [8]. Furthermore, the same research group showed a differential MAGI2 abundance in biopsies of patients with different glomerulopathies using immunofluorescence, revealing a marked downregulation of this protein in FSGS [8]. However, FSGS is a heterogeneous histological pattern for which several disease entities are responsible for. Especially the differentiation between primary (idiopathic FSGS without a known cause) and secondary FSGS (e.g. due to infections, maladaptation in obesity, drug toxicity) is still challenging. Until now, it is unclear whether MAGI2 is differentially expressed in these different FSGS subclasses and whether it could be a marker for the subclassification of this cohort.

To date, the search for new methods and new markers for the analysis of kidney biopsies to provide diagnoses and therapy for patients is proceeding quickly. In recent years, the use of modern microscopy techniques, such as super-resolution microscopy has improved the optical resolution of the filtration slit drastically [9]. 3D-structured illumination microscopy (3D-SIM), allows the morphometric analysis of the filtration barrier, and the filtration slit density can be determined by Podocyte Exact Morphology Measurement Procedure (PEMP) [5,10,11].

The zebrafish (*Danio rerio*) is a versatile animal model used in many different life science applications. Within the kidney field, zebrafish embryos have been used as a simple vertebrate model in which genetic modifications are rather easy with a conserved glomerular morphology and function [12] combined with a size small enough to even perform high-content drug screenings [13].

In zebrafish two orthologous forms, *magi2a* and *magi2b*, are reported for the human MAGI2 gene. Of these two orthologues, only *magi2a* has been shown to be expressed in the pronephric kidney, while *magi2b* was only expressed in the central nervous system [14]. In line with this finding, the knockout of *magi2a* but not *magi2b* resulted in slowly progressive proteinuria [14].

Herein, we investigated the expression of MAGI2 in murine glomerular disease models, isolated dedifferentiated murine glomeruli, isolated zebrafish glomeruli and patient biopsies using immunofluorescence analysis, tandem mass spectrometry, and mRNA sequencing. To investigate the co-regulatory relationship between Magi2 and Nephrin, we knocked out the respective proteins in larval zebrafish using CRISPR/Cas9 ribonucleoprotein (RNP) injection and observed the effects of protein absence on the development of proteinuria as well as on the expression of the respective other protein in the developing larvae.

## Results

### MAGI2 is expressed in healthy podocytes and localizes to the filtration slit

Independent of species (human, mouse, rat, zebrafish), we found that Magi2 was localized within the cell-cell contact of podocytes as well as colocalized with Nephrin (Figure 1 a-c) and the transgenic *nphs2*-driven mCherry reporter in podocytes of the zebrafish pronephros, respectively (Figure 1 d), as well the immunohistochemistry within the proteinatlas database (Suppl. Fig. 1A). In single-cell RNA sequencing datasets, *Magi2* clustered within the podocyte fraction, as shown in Supplemental Figure 1 B.

**Figure 1:**
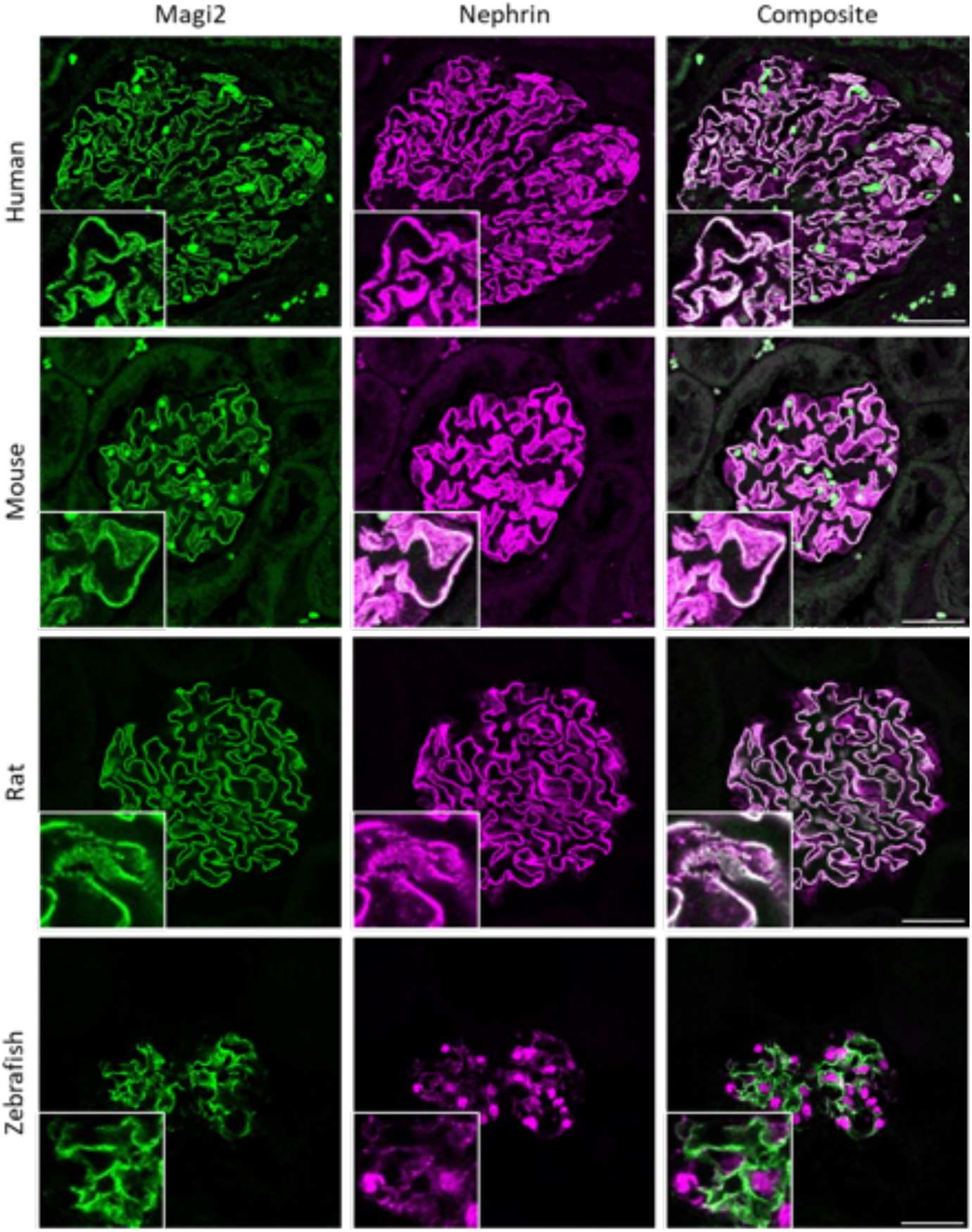
Confocal-LSM imaging of anti-MAGI2 stained kidney sections shows the binding of the MAGI2 antibody in human, mouse, rat kidneys and the zebrafish pronephros. Scale bars represent 20 µm.

To evaluate the subcellular localization of MAGI2 in podocytes *in situ*, we used super-resolution 3D-SIM of formalin-fixed paraffin-embedded (FFPE) tissue samples. The resolution obtained in this set of experiments was approximately 119±10 nm (mean±SD) as measured as the full width at half maximum (FWHM) of the subdiffractional slit diaphragm, which is sufficient to resolve individual podocyte foot processes in humans, rats, and mice. Super-resolution analysis of human kidney sections labeled for both NEPHRIN and MAGI2 revealed a direct localization of MAGI2 at the filtration slit demonstrated by colocalization with the slit diaphragm protein NEPHRIN (Figure 2).

**Figure 2:**
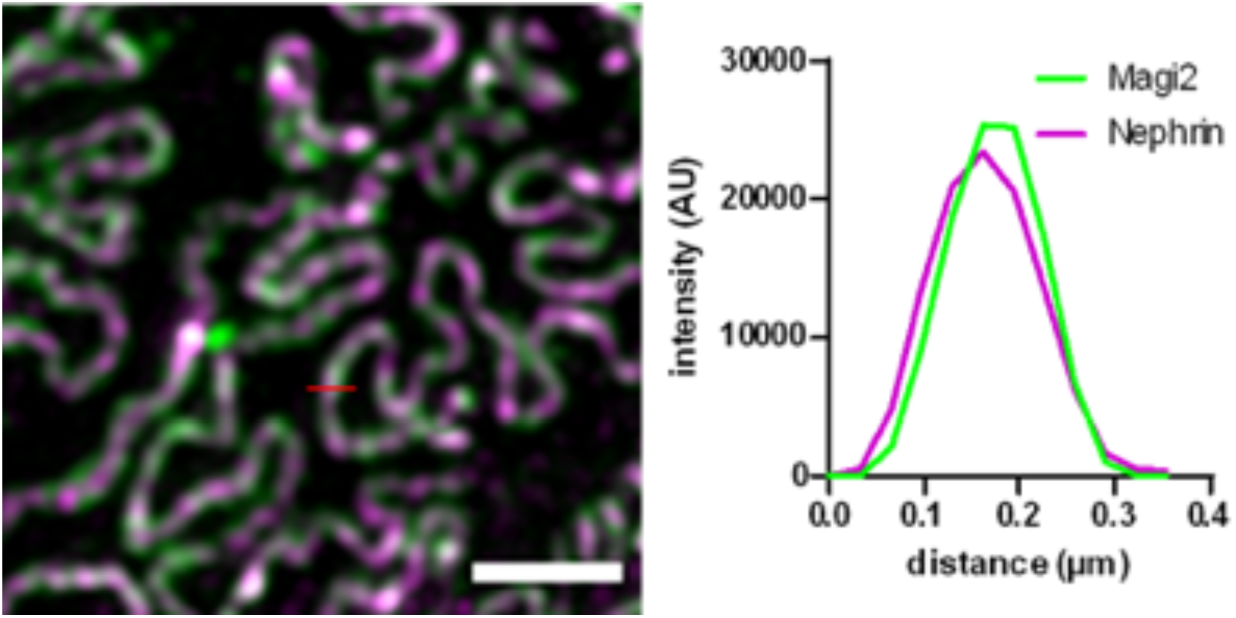
Super-resolution 3D-SIM microscopy of a human kidney section stained for MAGI2 (green) and nephrin (magenta) shows a colocalization of both proteins at the podocyte filtration slit. The scale bar represents 2 µm.

### MAGI2 is a sensitive marker for podocyte injury *in vitro* and *in vivo*

In the beforehand-established Glom*Assay* [15], glomeruli were isolated from C57BL/6 mice, and let dedifferentiate *in vitro* over time. In tandem mass spectrometric (LC-MS/MS) proteomic analysis generated from fresh and 6 days past isolation dedifferentiated glomeruli, Magi2 was statistically significantly downregulated accompanied by other podocyte-specific proteins like Alpha-actinin-4, Nephrin, Podocin, and Synaptopodin (Figure 3 A). In line with that, already three days after isolation, the Magi2 protein was significantly decreased in immunostained glomeruli (Figure 3 B). As shown in Figure 3 C, mRNA sequencing of glomeruli sampled freshly after isolation, as well as 3, 6, and 9 days of dedifferentiation, showed a progressive decrease of *Magi2* mRNA levels over the course of 9 days of cultivation and dedifferentiation. RT-qPCR analysis of glomeruli isolated freshly and after 6 days of cultivation confirmed a significant downregulation of *Magi2* (Figure 3 D).

**Figure 3:**
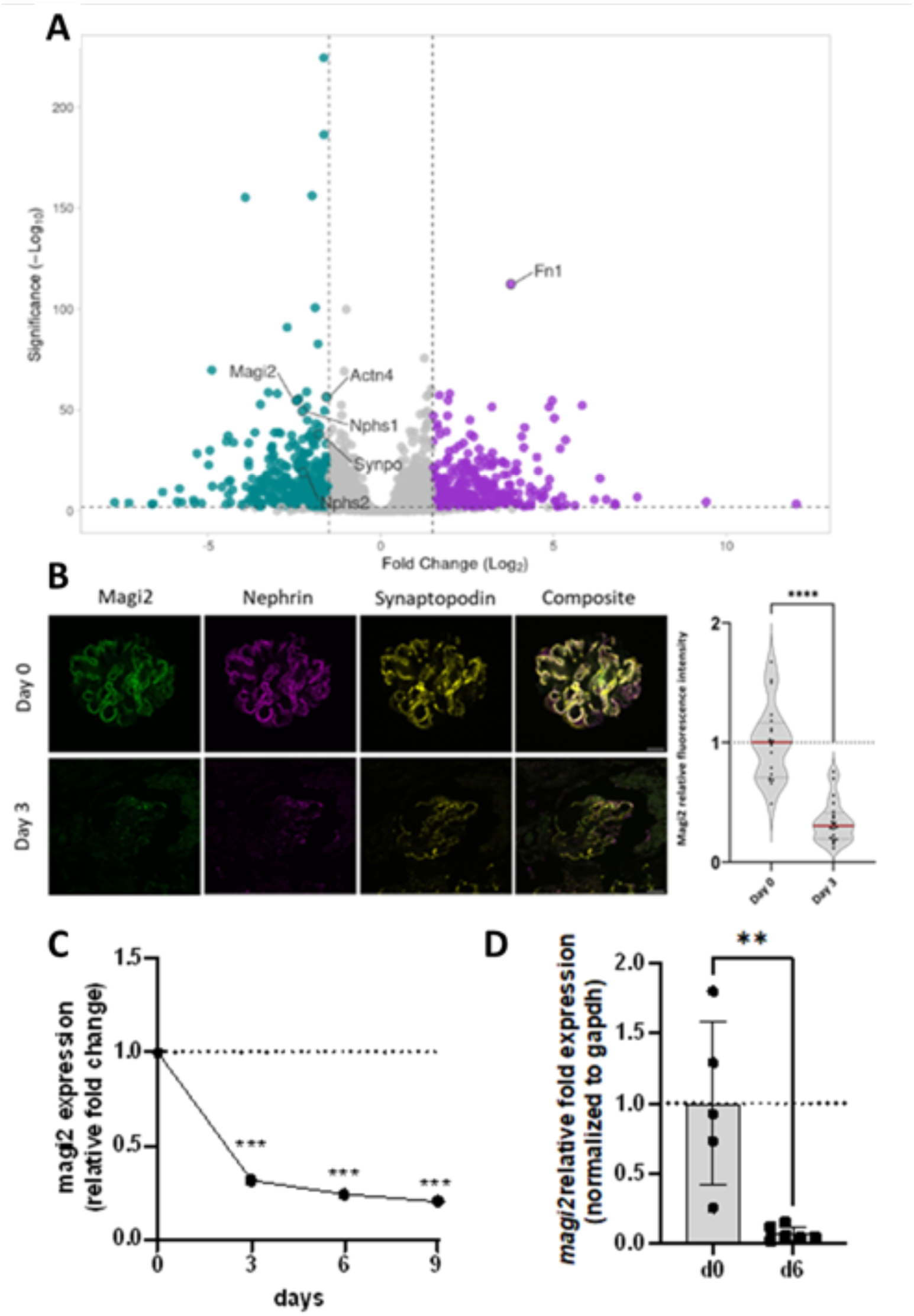
Time course of podocyte marker and MAGI2 in isolated cultured murine glomeruli. A shows the volcano plot of a proteomic analysis of dedifferentiated glomeruli at day 6 past isolation versus fresh isolation (GlomAssay). Magi2 and other relevant podocyte marker proteins like Nphs1, Nphs2, Synpo, and Actn4 were significantly downregulated while extracellular matrix proteins like Fn1 were increased. Immunofluorescence analysis for Magi2 and Nephrin verified this phenotype with a drastic increase of Magi2 already at day 3 past isolation. The scale bar represents 20 μm. Both mRNA sequencing (C) and RT-qPCR (D) showed significant downregulation of Magi2 mRNA starting from 3 days post isolation.

To rapidly evaluate the glomerular Magi2 protein regulation in animal models of glomerular diseases, we applied a Deep-Learning network (U-Net) that was trained to segment immunofluorescence-stained glomeruli in confocal laser scanning micrographs [16]. As shown in Suppl. Fig. 2, glomeruli were segmented from tissue sections with a Deep-Learning algorithm, and mean glomerular MAGI2 fluorescence intensity was analyzed in the resulting segmentation masks of the Synaptopodin-labeled glomeruli. In the uninephrectomy deoxycorticosterone acetate (DOCA) salt hypertension mouse model, mice develop hypertension, progressive albuminuria, and glomerulosclerosis after uninephrectomy, parenteral DOCA application, and oral high salt intake [17]. In this secondary FSGS model intraglomerular Magi2 fluorescence intensity showed a statistically significant decrease in DOCA mice compared to the uninephrectomized and sham-treated control animals (Figure 4 A). In the nephrotoxic serum nephritis (NTS) model, a murine model for crescentic glomerulonephritis, Magi2 was downregulated in comparison to control animals (Figure 4 B). Next, we evaluated glomerular Magi2 abundance in the puromycin aminonucleoside (PAN) nephropathy model. In this model, intravenous administration of PAN leads to reversible changes in podocytes, resulting in rapid proteinuria and foot process effacement mimicking minimal change nephropathy. In the PAN-injected animals, Magi2 was significantly downregulated (Figure 4 C).

**Figure 4:**
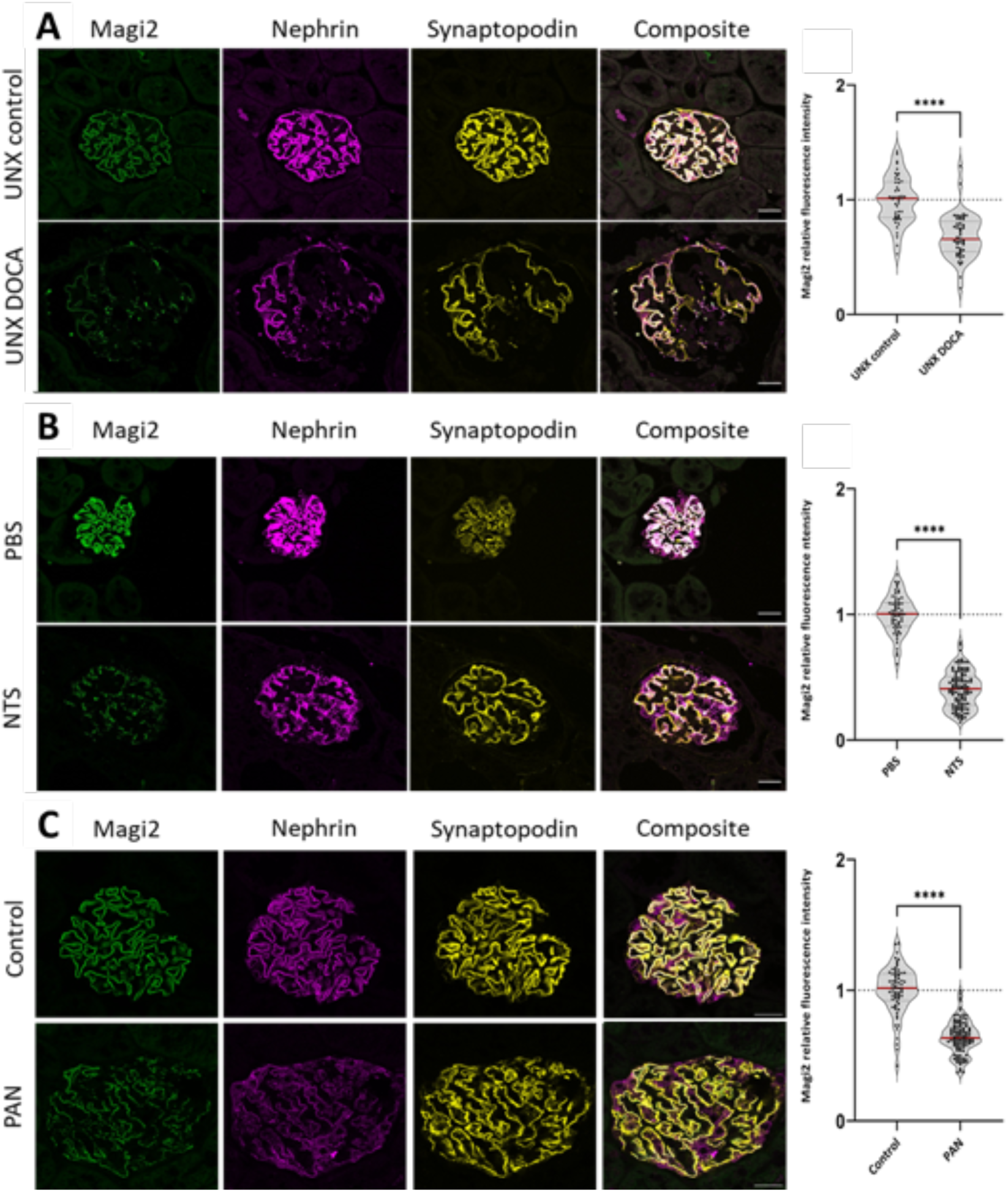
Magi2 protein was downregulated in the murine uninephrectomy DOCA salt hypertension model (A), the murine nephrotoxic serum nephritis (B) and the puromycin aminonucleoside nephropathy rat model (C). All scale bars represent 20 µm.

Having established that MAGI2 downregulation seems to be a stereotypical reaction upon podocyte injury, we wanted to elucidate whether this reaction is conserved across different vertebrate species. The podocyte-specific nitroreductase/metronidazole zebrafish model was used to induce FSGS-like disease in zebrafish upon subtotal podocyte depletion. In this model, podocytes in transgenic zebrafish larvae express a bacterial nitroreductase which intracellularly transforms MTZ to a cytotoxin [18]. This will lead to highly specific and dose-dependent ablation of podocytes. Larvae treated with 80 µM MTZ from 4-6 days post-fertilization (dpf) develop edema, proteinuria, and foot process effacement of the remaining hypertrophic podocytes [19]. In the acute phase of this model directly after induction of podocyte ablation (6 dpf), the glomerulus-specific zebrafish orthologue to MAGI2, Magi2a expression was significantly downregulated demonstrated by immunofluorescence stainings (Figure 5 A). As we have shown before, in the chronic phase after podocyte ablation, larvae develop FSGS-like disease including cellular lesions, activation of parietal epithelial cells, and deposition of extracellular lesions on the glomerular tuft[19]. Like the acute phase, Magi2a was significantly downregulated in the FSGS-like larvae as compared with the respective control animals (Figure 5 B). Additionally, we performed mRNA-sequencing of manually isolated glomeruli of larvae in the chronic phase (FSGS-like disease) and healthy controls which were treated with 0.1% DMSO. In this dataset, together with common podocyte targets like *nphs1*, *magi2a* was one of the top downregulated genes in the FSGS-like larvae in comparison to the control group (Figure 5 C).

**Figure 5:**
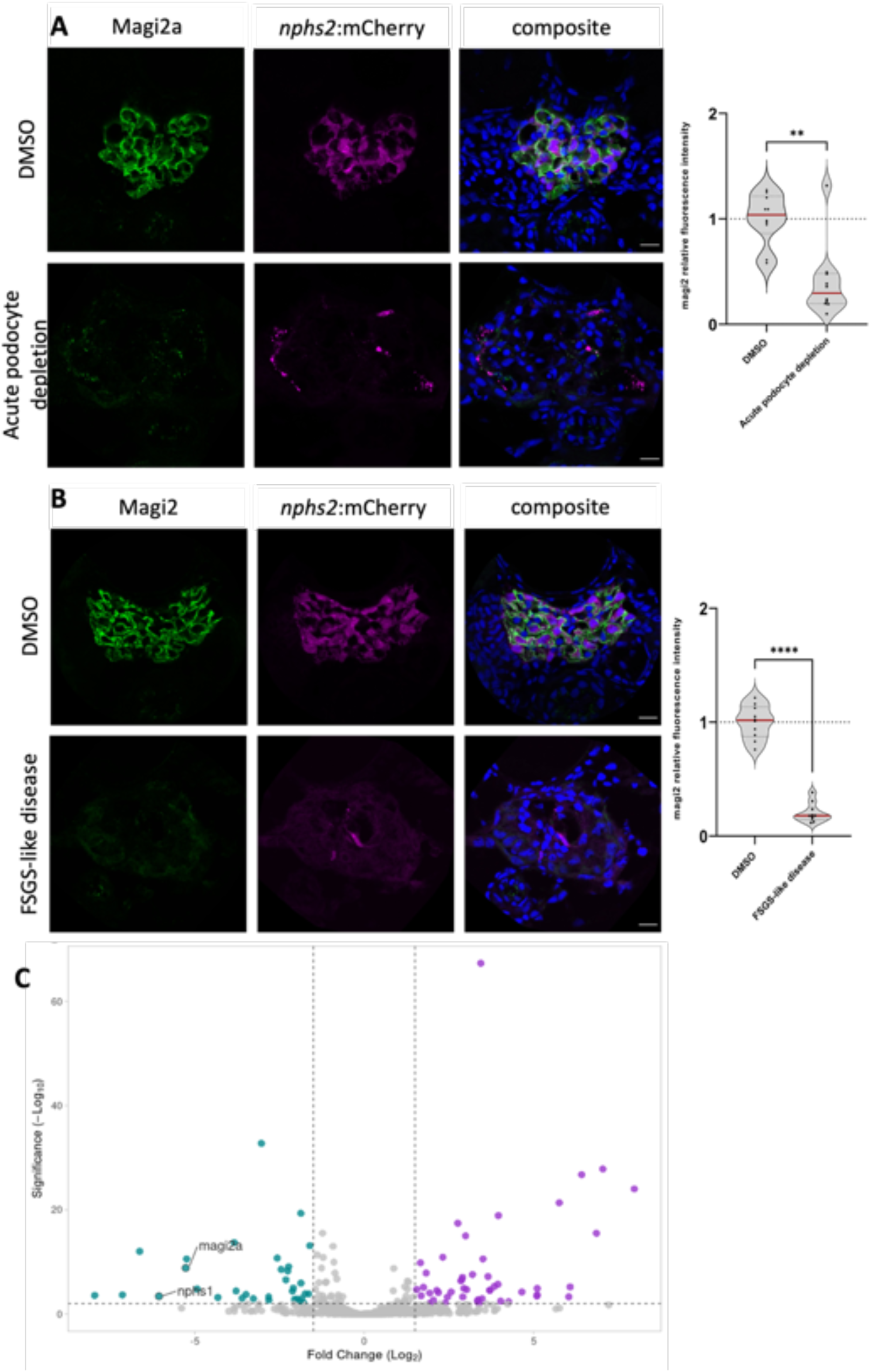
Downregulation of the zebrafish orthologue Magi2a was demonstrated both in the acute phase of pharmacogenetic podocyte depletion (A) as well as in the chronic FSGS-like phase (B). The volcano plot in C shows the downregulation of mRNA sequencing demonstrated statistically significant lower abundance of magi2a together with nphs1. Scale bars represent 20 µm.

### Generation of F0 generation zebrafish MAGI2 and NPHS1 podocytopathy models

Next, we created zebrafish CRISPR/Cas9 F0-mutants for *nphs1* and *magi2a* using injection of CRISPR/Cas9 ribonucleoprotein complexes (RNPs) targeting two different genomic regions within the *magi2a* or *nphs1* gene in one-cell stage embryos. Using this strategy, a high knockout efficiency can be achieved even in the first generation [20]. To rapidly screen for impairment of the glomerular filtration barrier, we used embryos of a transgenic strain expressing a vitamin D-binding protein eGFP fusion protein (gc-eGFP) in the blood plasma. Upon glomerular injury, this fusion protein will be filtered after loss of glomerular permselectivity. As a control, a guide RNA with no specific target within the zebrafish genome (targeting an intronic region in the human *HBB* gene) was used.

Immunofluorescence analysis for the Nephrin protein in *nphs1* and Magi2a in *magi2a* KO larvae showed a massive decrease in the glomerular abundance of the respective proteins indicating a high KO efficiency (Figure 6 A, B). Physiologically, the approximately 78 kDa gc-eGFP fusion protein does not traverse the glomerular filtration barrier in relevant amounts and therefore serves as a sensitive marker for GFB impairment and proteinuria. In the immunostainings, we noticed eGFP-positive vesicles within proximal tubule cells which is indicative of glomerular barrier function defects (Figure 6 A, B). Additionally, using *in vivo* fluorescence microscopy, we found eGFP in the tubules of both *nphs1* and *magi2a KO* larvae. In contrast to that, the control-injected larvae did not show any eGFP vesicles in the proximal tubules. To analyze whether the KO leads to relevant clearance of high molecular weight protein from the blood, we used an automated high-content screening approach to quantify intravascular fluorescence intensities in a large number of larvae, which was established before [13]. In both mutant lines, a significant reduction of gc-eGFP fluorescence in the vasculature was observed. However, while the *nphs1* knockout causes a decrease of eGFP fluorescence as early as 4 dpf, the earliest difference in the *magi2a* KO larvae was evident at 5 dpf (Figure 7). The ultrastructural evaluation of these larvae revealed a full and diffuse podocyte foot process effacement in both models with preserved morphology of the glomerular basement membrane and glomerular endothelial cells (Figure 8).

**Figure 6:**
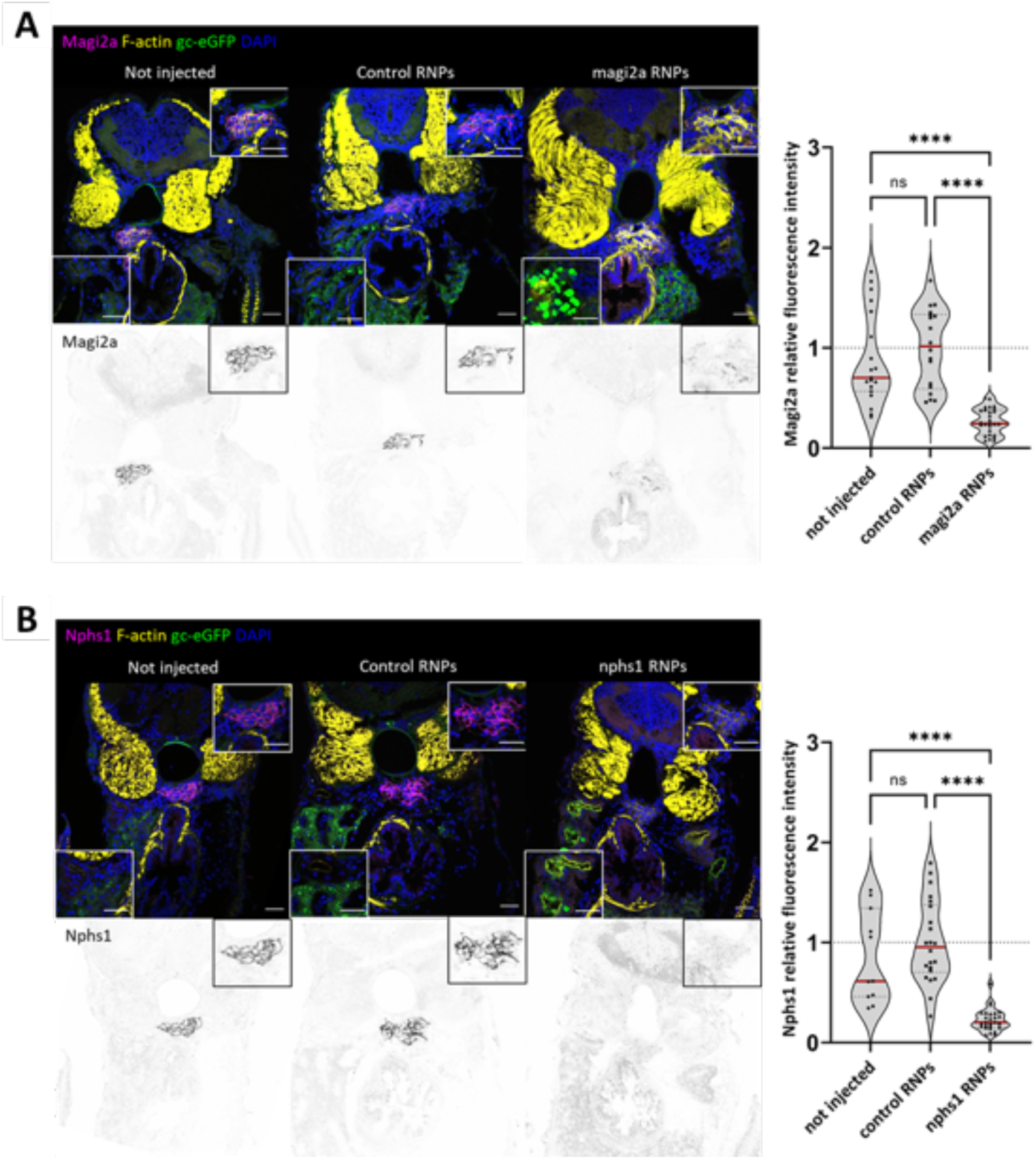
Generation of magi2a and nphs1 CRISPR/Cas9 mutants. A shows the decrease of glomerular Magi2 protein levels was observed only in the magi2a RNP, but not the control or uninjected embryos. B shows the massive decrease of Nephrin protein expression on the nphs1 mutant embryos. All scale bars represent 10 µm.

**Figure 7:**
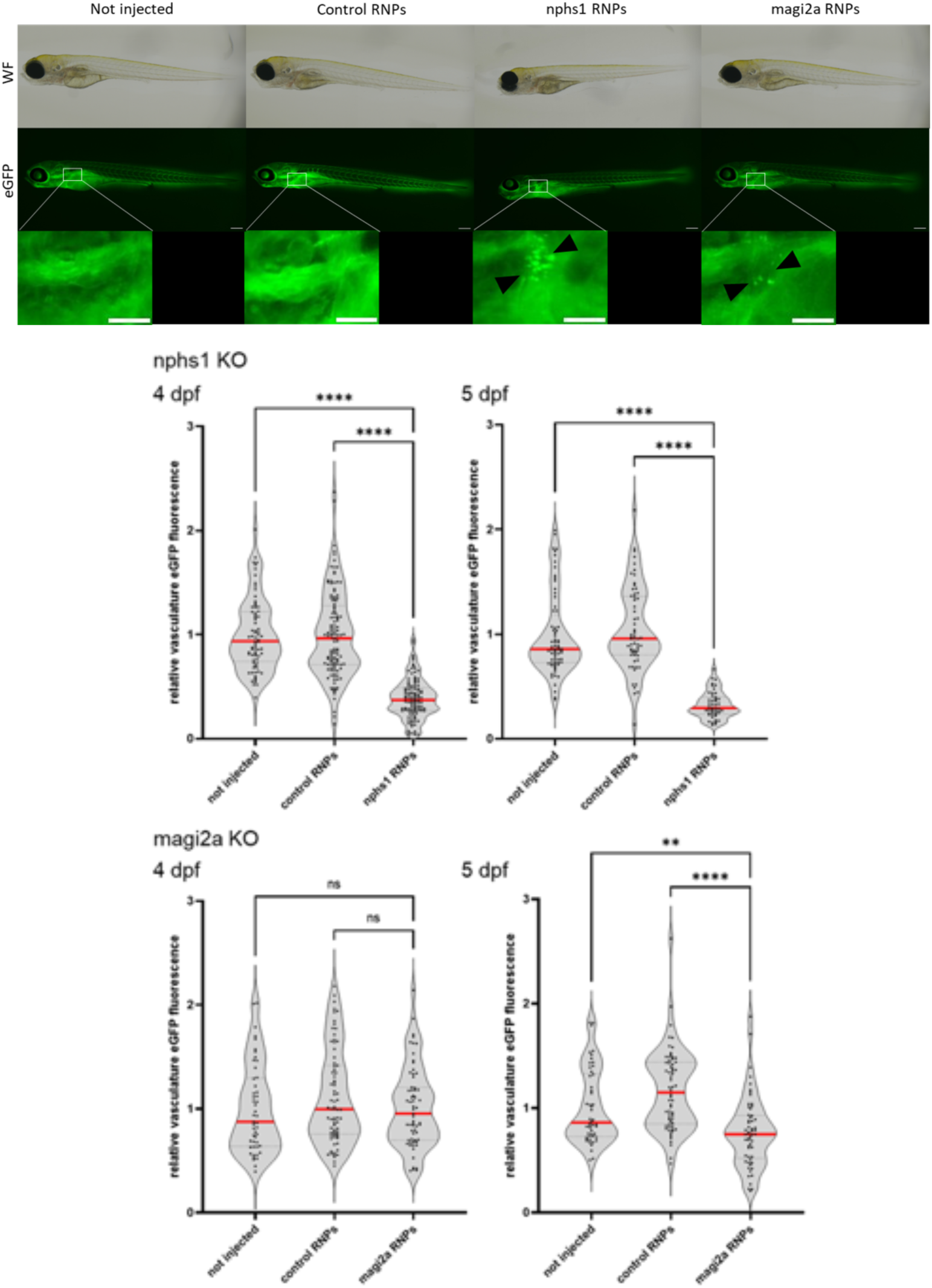
Evaluation of proteinuria in nphs1 and magi2a KO embryos. A shows the general morphology of larvae which were either uninjected, injected with control RNPs, or with nphs1 or magi2a RNPs. The fluorescence microcraphs in B show the filtered and reabsorbed gc-eGFP fusion protein in proximal tubules, but not in control- or uninjected embryos. In the beforehand established high-content proteinuria screening approach, nphs1 KO embryos were proteinuric at day 4 and 5 as demonstrated by decreased intravascular gc-eGFP fluorescence, while magi2a KO embryos were significantly proteinuric later at day 5. Scale bars indicate 200 µm in the overview and 100 µm in the zoom inserts.

**Figure 8.**
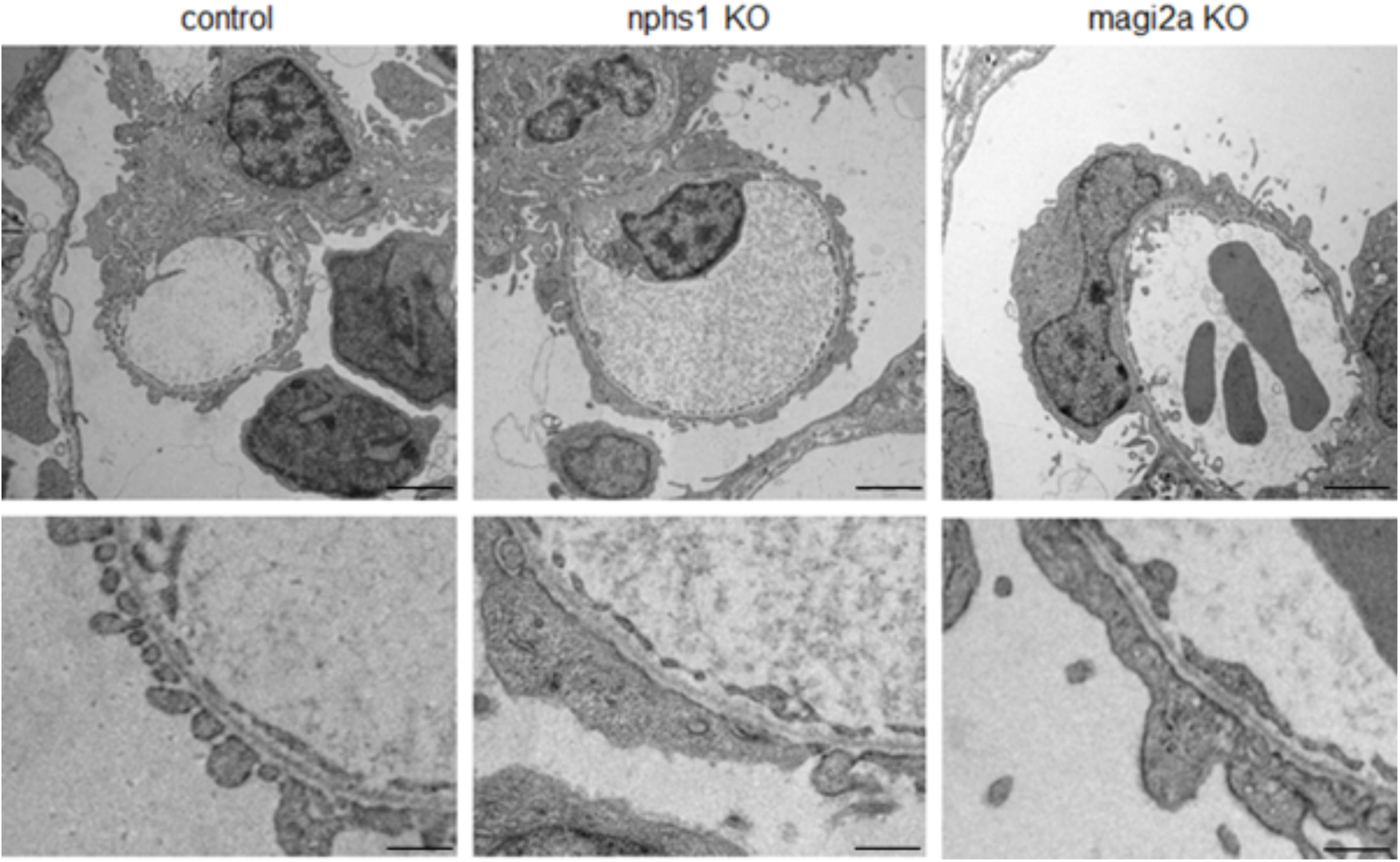
Ultrastructural analysis of control, nphs1 and magi2a KO embryos at 4 dpf revealed diffuse foot process effacement in both nphs1 and magi2a KO larvae with unaltered morphology in the other cell types or the extracellular matrix. Scale bars represent 2 µm in the upper and 500 nm in the lower panel.

### Coregulation of Magi2a and Nphs1 in genetic podocytopathy zebrafish models

As shown in Figure 9, immunofluorescence analysis of the *magi2a* KO but not the control-injected larvae showed a statistically significant decrease of Nephrin. Interestingly, vice versa in the *nphs1* KO larvae the expression of Magi2a was not changed.

**Figure 9.**
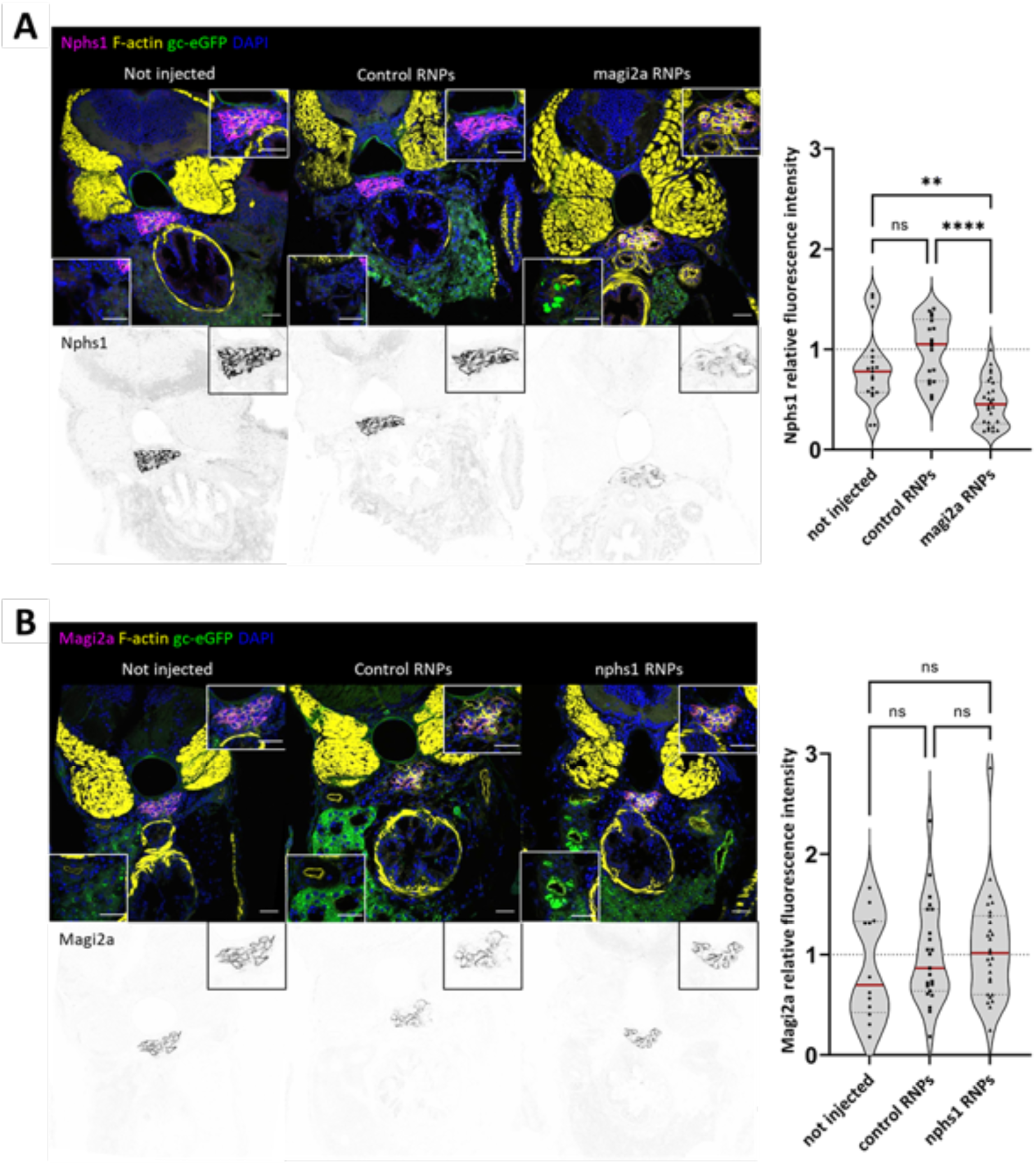
magi2a is required for the correct localization of nphs1 but not vice-versa. Immunofluorescence analysis in magi2a KO, but not control-injected larvae, showed a decrease of nphs1 protein (A). Vice-versa, KO of nphs1 had no influence on the localization of magi2a (B). Scale bars represent 10 µm.

### MAGI2 as a marker for primary FSGS

Having established that MAGI2 downregulation seems to be a stereotypical reaction in glomerular disease models, we evaluated its protein level in immunofluorescence analysis as a possible marker for glomerular diseases (Figure 10 A). We analyzed stained archived FFPE kidney biopsies of patients affected by MCD and FSGS classified as either primary or secondary. Proteinuria and serum creatinine as well as info about the presence of nephrotic syndrome is summarized in Table 1.

**Figure 10:**
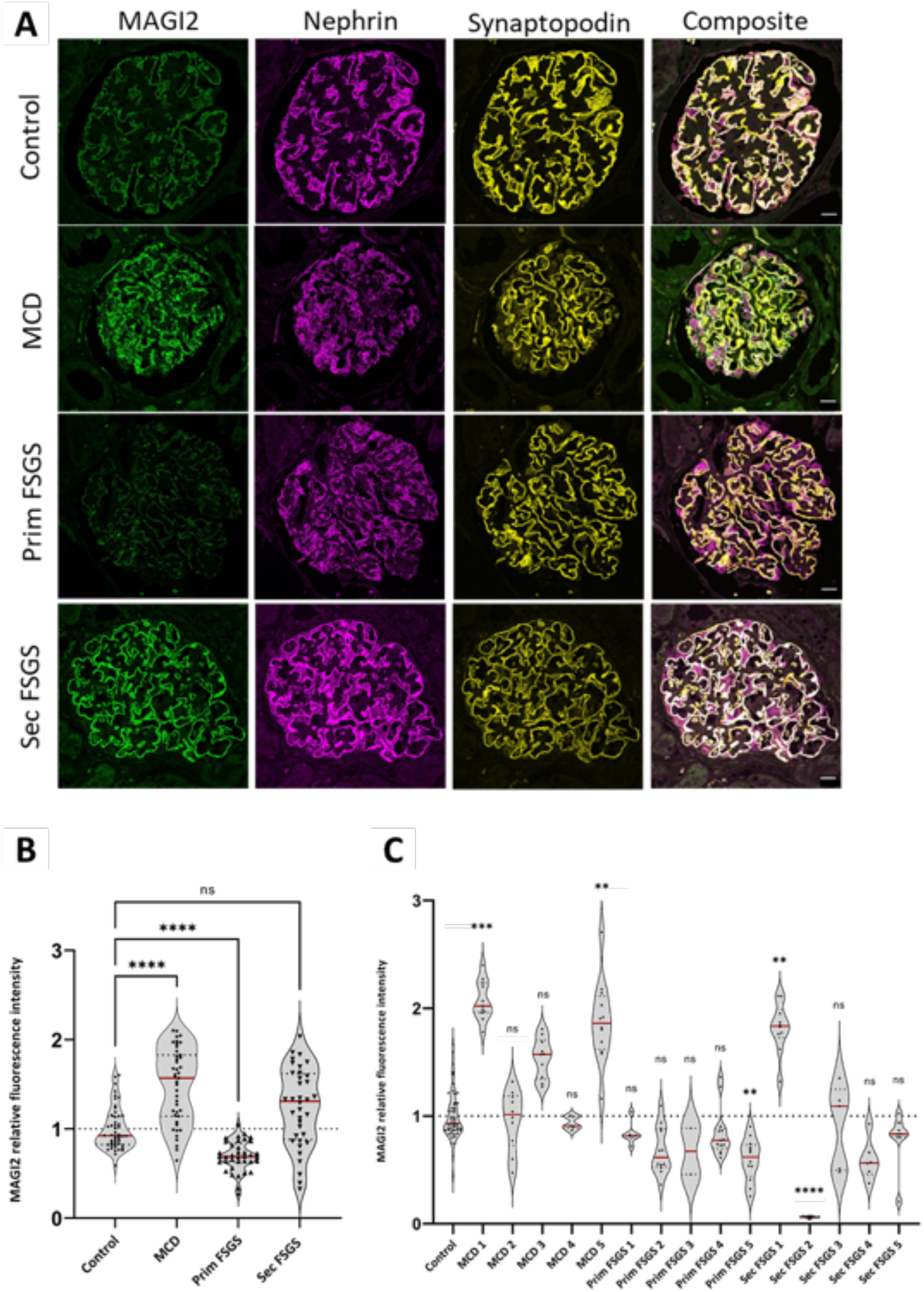
Glomerular MAGI2 expression is differentially regulated in MCD, primary FSGS, and secondary FSGS. In comparison to the healthy control nephrectomy tissue. Over all patient biopsies, (B) MAGI2 was significantly increase in the MCD group, while it was decreased in the primary FSGS, and unaltered in the secondary FSGS group. The scale bars represent 20 µm.

**Table 1:** Patient characteristics of the patients of which kidney biopsies have been available for analysis.

| Samples | Proteinuria | Creatinine | Nephrotic syndrome |
| --- | --- | --- | --- |
| MC 1 | 15.7 g/g creatinine | 2,12 mg/dl | yes |
| MC 2 | 10 g/g creatinine | 0.72 mg/dl | yes |
| MC 3 | 5 g/l | 0,59 mg/dl | yes |
| MC 4 | >0.5 g/dl | 3,5 mg/dl | yes |
| MC 5 | 9,7 g/24h | 1,2 mg/dl | yes |
| Prim FSGS 1 | 5 g/24h | 1,51 mg/dl | n.d. |
| Prim FSGS 2 | 9 g/24h | 1.28 mg/dl | yes |
| Prim FSGS 3 | 6 g/24h | 1.11 mg/dl | yes |
| Prim FSGS 4 | 8,4 g/g creatinine | 0,7 mg/dl | yes |
| Prim FSGS 5 | 6.08 g/24h | 0,7 mg/dl | yes |
| Sec FSGS 1 | 2.3 g/24h | 1,2 mg/dl | no |
| Sec FSGS 2 | 10.8 g/g creatinine | 4,14 mg/dl | no |
| Sec FSGS 3 | 4 g/24h | 2,3 mg/dl | no |
| Sec FSGS 4 | >8 g/24h | 4,67 mg/dl | no |
| Sec FSGS 5 | 4.8 g/24h | 3,19 mg/dl | no |

Interestingly, we found differential regulation of this protein between the groups analyzed: Compared to healthy controls, we observed that MAGI2 abundance was increased in 2/5 patients diagnosed with MCD, while a statistically significant downregulation could be demonstrated in cases of FSGS (Figure 10 B). Interestingly, when looking at the FSGS subgroups, this downregulation was limited to cases of primary FSGS, while the expression of MAGI2 in cases of secondary FSGS was unchanged (Figure 10 C).

Tight junction components of the filtration slit in podocytes can change their localization upon injury and podocyte foot process effacement [5]. As exemplarily shown in Figure 11, MAGI2 was localized in both healthy-appearing filtration slits (dense filtration slits meandering on the capillary) and effaced areas (linearized filtration slits) still colocalizing with NEPHRIN.

**Figure 11:**
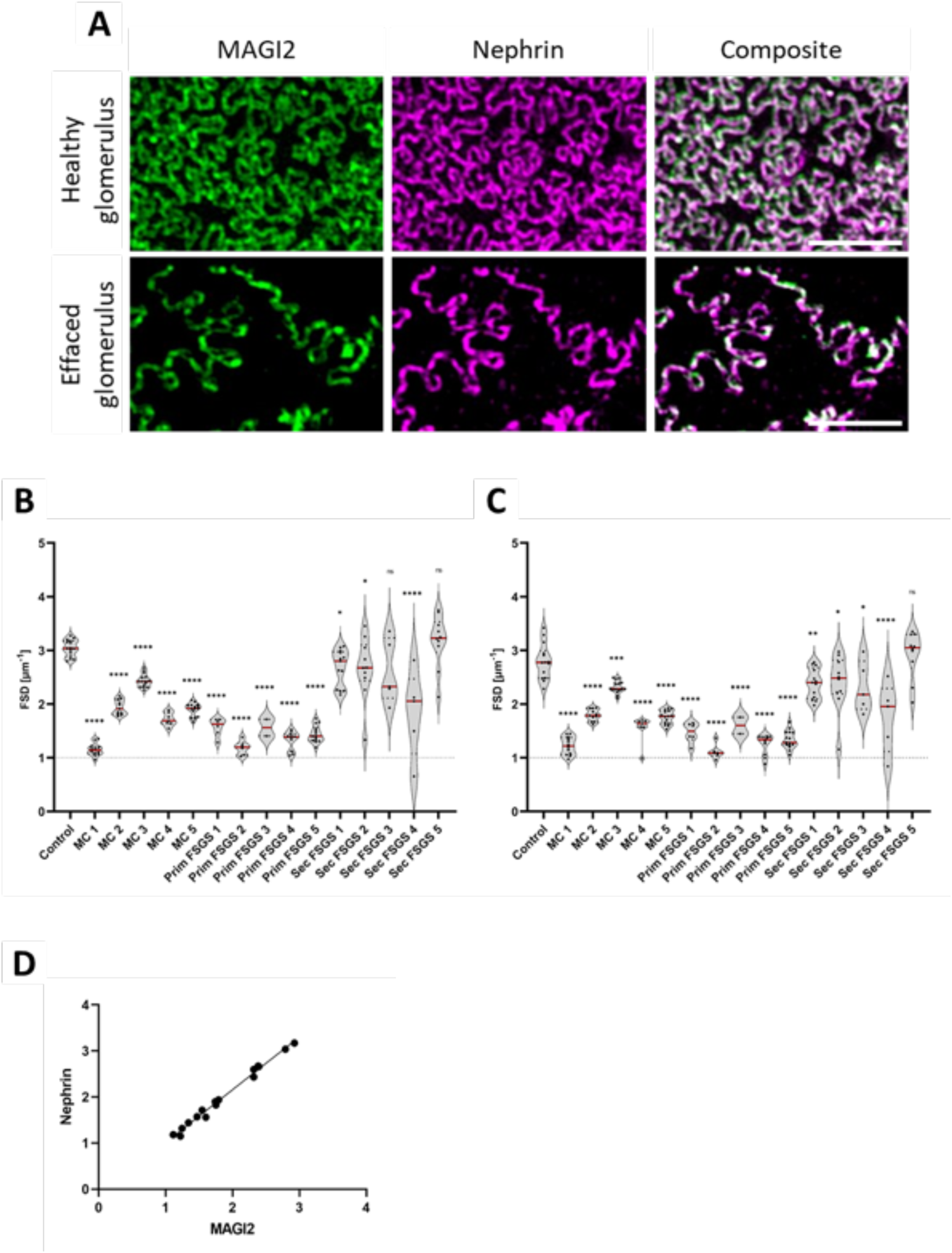
The use of MAGI2 as a marker for super-resolution microscopy enabled morphometry. As shown in (A), MAGI2 was co-localized with NEPHRIN both in healthy and in effaced podocytes. Both MAGI2 (B) and NEPHRIN (C) were used as a marker protein to determine the filtration slit density as a marker for foot process effacement. The graph in (D) shows the strong positive correlation of both values when determined for the same biopsy. The scale bars represent 2 µm.

NEPHRIN is regularly used as a marker for the calculation of filtration slit density (FSD) through the established Podocyte Exact Morphology Measurement Procedure (PEMP)[11]. To investigate whether MAGI2 could be used as a marker for glomerular filtration barrier disruption in PEMP, FSD was measured on human biopsies by PEMP with MAGI2 (Figure 11 B) or NEPHRIN (Figure 11 C). Both values showed a highly significant positive correlation (Figure 11 C) indicating interchangeability of both proteins for the determination of foot process effacement.

## Discussion

Chronic kidney disease (CKD) is a global public health burden affecting over 500 million people worldwide [21]. Podocytopathies and the associated impairment of the glomerular filtration barrier (GFB) are responsible for approximately 70% of all cases of CKD [3]. Although considerable progress has been made in recent years in understanding the molecular mechanisms involved in the onset of these diseases, much remains to be discovered.

The critical function of the MAGI2 protein and its podocyte-specific role has been investigated using *Magi2* knockout mice [8,22]. MAGI2 is of fundamental importance for both the formation and maintenance of the GFB, thus preventing the disruption of the slit diaphragm and the morphological abnormalities of the foot processes.

In the kidney, MAGI2 is expressed exclusively in the glomerulus, and within the glomerulus specifically in podocytes. Herein, we performed the first super-resolution microscopy study on the MAGI2 expression in human biopsies. 3D-SIM revealed that MAGI2 localizes at the filtration slit and directly colocalizes with the slit diaphragm protein Nephrin in all species investigated. Speaking for the zebrafish model, these results once again prove the high homology of the glomerular filtration barrier between vertebrate species in general, and the high value of this model as it has been demonstrated before [23].

The proposed implication of MAGI2 in glomerulopathies is supported by both animal and human studies. In lipopolysaccharide (LPS) treated mice, a strong reduction of *Magi2* gene expression coincided with podocyte injury [24]. In a rat model of crescentic glomerulonephritis (CGN) the downregulation of Magi2 was associated with an accumulation of Dendrin in the nucleus of podocytes [8]. This result, together with the knowledge that nuclear relocation of Dendrin in response to glomerular damage promotes podocytes apoptosis, suggests for Magi2 a role in avoiding podocyte death by sequestering Dendrin at the SD level and preventing its nuclear translocation [25]. In humans, recessive mutations in the *MAGI2* gene lead to steroid-resistant nephrotic syndrome [7,26].

To further evaluate the regulation of MAGI2 in glomerular diseases, we included in our study four different glomerulopathy animal models: The DOCA-salt hypertension heminephrectomy mouse model, the nephrotoxic serum nephritis mouse model, the puromycin aminonucleoside model, and zebrafish genetic podocytopathy models.

Measurement of the fluorescence intensity of Magi2 showed a significant reduction in the expression of this protein in DOCA and NTS treated mice, which mimic human secondary FSGS and crescentic glomerulonephritis, respectively. Similar results were found for the PAN nephropathy rat model which is a model for human MCD.

The zebrafish genome features genome duplications, so that two orthologous forms, *magi2a* and *magi2b*, are reported for the human *MAGI2* gene in zebrafish. The expression of *magi2*, without distinction between the two orthologous forms, has been previously demonstrated in the pronephric zebrafish glomeruli [27]. Subsequently, it was shown that *magi2a* is the only functional zebrafish orthologue of human MAGI2 in the pronephros [14]. We demonstrated the conserved localization of Magi2a protein in the slit diaphragm of zebrafish larvae and its colocalization with Nphs1. Analogue to the mammalian response to glomerular injury, Magi2a was downregulated both in the acute and the chronic phase of injury in this model.

Since the dedifferentiation of podocytes plays a key role in the development of different types of glomerulopathies we wondered whether the regulation of Magi2 in the investigated models is a stereotypical consequence of podocyte damage in general or occurs due to podocyte dedifferentiation. In the established *GlomAssay* [15], glomeruli are isolated from mice and dedifferentiated *in vitro* over time. Herein we show a statistically significant downregulation of podocyte-specific proteins, and among them slit diaphragm proteins like Nephrin, Podocin, Cd2ap and Magi2 were the most strongly down-regulated.

Furthermore, it has been shown that, as demonstrated for other scaffold proteins such as CD2AP [28], ZO-1 [29] and NEPHRIN [30] itself, the expression of MAGI2 is reduced in glomerulopathies. Using different glomerulopathy animal models has allowed us to verify that the results are independent of the species and the model investigated. The regulation of MAGI2 in glomerulopathies appears to be a consequence of the process of podocyte dedifferentiation.

In humans, a different regulation of MAGI2 in different glomerulopathies and, in particular, a marked downregulation of this protein in FSGS has been reported [31]. FSGS is a heterogeneous histological pattern in which several subgroups are included: primary, secondary, and genetic FSGS. MAGI2 expression was slightly increased in biopsies of patients diagnosed with MCD. Looking at all biopsies investigated, a statistically significant mean downregulation could be demonstrated in cases of primary FSGS but not in cases of secondary FSGS. From the correlation between the clinical data of the patients whose biopsies were investigated, and the results obtained from the measurement of the fluorescence intensity of MAGI2, there was no direct correlation between the development of nephrotic syndrome and the downregulation of this protein, suggesting that other proteins and other mechanisms in addition to the reduced expression of MAGI2 may have a role in the onset of the nephrotic syndrome. Using super-resolution 3D-SIM we have previously shown that slit diaphragm components, such as CLDN5, can change their localization in case of glomerulopathy. Differently, MAGI2 continues to colocalize with NEPHRIN even in effaced areas, and like NEPHRIN can be used as a marker for the calculation of filtration slit density (FSD) using the PEMP [11].

The interaction of MAGI2 with other SD proteins, such as NEPHRIN and DENDRIN, has been implicated in numerous biological processes, including the regulation of apoptosis, SD protein expression, and podocyte cytoskeletal reorganization [22]. Although several studies have demonstrated a physical interaction between Magi2 and Nephrin [4,24], suggesting the possibility that Magi2 orchestrates the localization of Nephrin in the podocyte slit diaphragm [31], there are still pros and cons regarding the interaction between these two proteins. In particular, two studies reported the interaction of Magi1 (another member of the MAGUK family) but not Magi2 with Nephrin [32,33].

We created CRISPR / Cas9 mutants for the zebrafish orthologues of *nphs1* and *magi2a*. First, we evaluated the development of proteinuria in both mutants. We have shown the presence of eGFP-positive vesicles within proximal tubules following knockout of both genes, with *in vivo* fluorescence microscopy and with immunostaining of cryosections of larvae at 5 days post fertilization (dpf). The glomerular clearance of gc-eGFP leads to a reduction in intravascular fluorescence, which can be calculated using an innovative and automated high-contest screening approach [13]. We observed that the *nphs1* KO caused a reduction in the fluorescence of eGFP already at 4 dpf, while in the *magi2a* KO larvae the earliest difference was evident at 5 dpf. The premature appearance of proteinuria in zebrafish larvae KO for *nphs1* compared to larvae ko for *magi2a* would confirm that the presence of proteinuria following the absence of Magi2a is a consequence of the downregulation of Nephrin. Interestingly, in *magi2a* knockout larvae we found a statistically significant decrease in the glomerular abundance of Nphs1, while in the *nphs1* KO larvae the expression of Magi2a was not changed. Although these results do not allow us to demonstrate a physical interaction between Nphs1 and Magi2a, they still suggest a relation between the two proteins, and in line with Yamada and colleagues, they confirm the hypothesis that the presence of Magi2 is fundamental for the localization of Nephrin.

To summarize, we have shown that the expression of MAGI2 in the glomerulus is associated with the expression of nephrin regardless of the species investigated (human, mouse, rat, zebrafish) and that its expression is reduced in animal models of glomerulopathies. We showed that Nephrin localization within the slit diaphragm is dependent on Magi2a and not vice versa emphasizing its critical role as a scaffold protein. Due to the specific regulation of MAGI2 depending on the podocytopathy investigated, we propose a role for MAGI2 to distinguish between FSGS and MCD and its use as a marker for disruption of the glomerular filtration barrier.

## Material and Methods

### Kidney Biopsy material

For our study formalin-fixed paraffin-embedded (FFPE) human kidney biopsies of patients diagnosed with MCD (n = 5), primary FSGS (n=5), and secondary FSGS (n = 5) by experienced pathologists of Cologne (part of the FOrMe-registry (NCT03949972), a nationwide registry for MCD and FSGS in Germany) were used. As healthy controls, anonymized excess normal kidney tissue of partial nephrectomies of the Department of Urology of the University Medicine Greifswald was used (n = 5). The use of excess kidney tissue from tumor nephrectomies has been approved by the Ethics Committee of the University Medicine Greifswald (Ref. No. BB 075/14). All patients stated written informed consent. All experiments were performed in accordance with local guidelines overseen by the University Medicine Greifswald and University Greifswald, Greifswald, Mecklenburg— Western Pomerania. Four-micrometer sections were cut and mounted on Superfrost slides (R. Langenbrinck GmbH).

FFPE kidney tissue of the uninephrectomy DOCA salt hypertension and of nephrotoxic serum (NTS) mouse models and of the PAN treated rat model of previous studies was used, which was prepared as described before [17].

### Glomerulus isolation

Murine glomeruli of male 6 months old nphs1:CFP mice were isolated as described before[15]. Directly after isolation and after 3 days of dedifferentiation, glomeruli were centrifuged at 3500 rpm for 3 min and fixed in 2% PFA for 10 min. After two times washing in PBS, they were embedded in molten 2% agarose at 37°C. After hardening at room temperature, agarose blocks were left overnight in ethanol and embedded in paraffin using the standard routine for kidney tissue.

### Proteomic analysis of isolated, dedifferentiated glomeruli

Murine glomeruli of male 6 months old C57Bl/6J mice were isolated as described above. Glomeruli were snap-frozen directly after isolation and after 6 days of dedifferentiation and subjected to LC-MS/MS analysis as described before. Briefly, the samples were subjected to tryptic digestion and analyzed by tandem mass spectrometry in data-independent mode on an Ultimate 3000 - Exploris 480 LC-MS configuration. The dataset is made publicly available on the PRIDE database under the accession number PXD056696.

### RNA-isolation and RT-qPCR from isolated glomeruli

Total RNA was isolated using the TRIreagent (Sigma Aldrich) and reverse transcribed with the Qiagen reverse transcription kit as described before[34]. A duplex Taqman RT-qPCR was performed using FAM-conjugated probes targeting Magi2 (Mm01159076_m1) and VIC-conjugated probes against Gapdh (Mm99999915_g1) both produced by ThermoFisher Scientific with the Luna Universal Probe qPCR Master Mix (NEB, 3004L) on the QuantStudio 3 Real-Time PCR System (ThermoFisher Scientific). Amplification data were analyzed within the QuantStudio software package with the ΔΔCt method with Gapdh as a reference gene and normalized as relative fold expression to freshly isolated glomeruli (day 0). Statistical difference was evaluated in GraphPad Prism 9 (V 9.3.1)

### Immunofluorescence staining

For mouse, rat and human sections and for murine isolated glomeruli, after deparaffinization in xylene and rehydration in a descending ethanol series, all sections were boiled with a pressure cooker in Tris EDTA buffer containing 0.1% Tween-20 (10 mmol/L Tris, 1 mmol/L EDTA, pH = 9) for antigen retrieval. After autofluorescence quenching in 100 mmol/L glycine diluted in purified water and three times washing in PBS, slides were incubated for 1 hour in blocking solution containing 1% fetal bovine serum, 1% normal goat serum, 1% cold fish gelatine, and 1% bovine serum albumin. The primary antibodies (1:150 in blocking solution, polyclonal rabbit anti-MAGI2 IgG, HPA013650, Sigma; 1:100 in blocking solution, polyclonal Guinea pig anti-Nephrin IgG, GP-N2, Progen; 1:50 in blocking solution, monoclonal mouse anti-Synaptopodin IgG, 61094, Progen) were incubated at 4°C on the slides overnight. The next day after five times washing in PBS and blocking for one hour, the secondary antibodies (1:800 in blocking solution Cy3-conjugated donkey anti-guinea pig IgG (H+L), 706-165-148, Jackson Immuno Research, Hamburg, Germany; 1:2000 Nano-Secondary® alpaca anti-human IgG/anti-rabbit IgG, recombinant VHH, Alexa Fluor® 488 [CTK0101, CTK0102]; 1:2000 Nano-Secondary® alpaca anti-human IgG/anti-rabbit IgG, recombinant VHH, Alexa Fluor® 647 [CTK0101, CTK0102]) were incubated at room temperature for one hour, followed by five times washing in PBS and transfer to purified water. The slices were mounted in Mowiol for Microscopy (Carl Roth) using High Precision cover glasses (Paul Marienfeld GmbH).

The zebrafish sections were permeabilized with 0.1% Triton X-100 in PBS. Antigenic epitopes were demasked with 1% SDS and the sections were then incubated for one hour in blocking solution. The sections were incubated with respective primary antibodies (1:1000 in blocking solution, polyclonal rabbit anti-MAGI2 IgG, HPA013650, Sigma; 1:2000 rabbit anti-zebrafish nephrin, Innovagen, Lund, Sweden) at 4 °C overnight. After three washes in 1x PBS and incubation with the Alexa 647 conjugated goat anti-rabbit F(ab) antibody fragment (1:800) the slides were incubated with 0.013mg/ml Hoechst 33342 (Sigma-Aldrich) for 10 minutes, washed in 1xPBS and mounted in Mowiol for microscopy.

### Fluorescence microscopy

3D-SIM microscopy was performed as described before with slight differences in the reconstruction parameters: Baseline Cut, SR Frequency Weighting: 1.0; Noise Filter: −5.6; Sectioning: 97, 84, 84[11]. For confocal laser scanning microscopy, a Leica TCS SP5 (Leica Microsystems) and an OLYMPUS FV3000, both equipped with a 40 × (NA 1.4) oil immersion objective, were used. Mouse glomeruli were imaged with 3.9691 pixels per micron, human glomeruli were imaged with 2.64 pixels per micron and zebrafish glomeruli were imaged with 9.6544 pixels per micron. Images were exported as .czi and .oir files.

### Deep-learning-enabled quantitative image analysis

A custom-developed FIJI script was used to segment glomeruli in kidney cross-sections using a custom-trained Deep-Learning network. To enable this, a UNet was trained within the ZeroCostDL4Mic ecosystem[35] which has been published before[16]. The resulting network was imported to FIJI using DeepImageJ. The resulting predictions from the network were used to generate regions of interest (ROIs) in which mean fluorescence intensity was quantified over all channels. The FIJI macro script can be accessed at http://github.com/Siegerist.

### Zebrafish experiments

Zebrafish were bred as described before. All zebrafish experiments adhered to national law and local ethical standards as overseen by the “Landesamt für Landwirtschaft, Lebensmittelsicherheit und Fischerei Mecklenburg-Vorpommern”. The zebrafish FSGS-like disease model was used as described before[19]. Herein, zebrafish larvae transgenically express the bacterial enzyme nitroreductase and the fluorescent protein mCherry exclusively in podocytes. Treating them with a concentration of 80 µmol L − 1 of metronidazole (MTZ) will lead to partial podocyte depletion and development of FSGS-like lesions. Treatment started at 4 dpf, lasted 48 hours and the larvae were collected for histological analysis at 6dpf and 8dpf.

### Zebrafish glomerulus isolation and mRNA sequencing

Larvae were anesthetized with Tricaine and collected in a tube, E3 medium was discarded, L15 medium and 20 ceramic beads were added. The homogenization was performed in a tissue homogenizer. The lysed sample was resuspended in L15 medium and transferred to a petri dish. mCherry-positive glomeruli in lysed samples were collected using a P100 micropipette under the fluorescent microscope NIKON SMZ18. The isolated glomeruli were transferred in a tube and collected by centrifugation at 1000 xg. The pellet was stored at −80°C. RNA isolation was performed using the miRNeasy Micro Kit from Qiagen according to the manufacturer’s protocol. The sequencing was performed by GenXPro (Frankfurt, Germany). MACE-Seq for ultra-low input was used for the mRNA-Seq library preparation. mRNA Seq library was used to perform sequencing on control and MTZ samples.

### Zebrafish CRISPR/Cas9 knockout

crRNAs and Atto-550 tracrRNA were designed using ChopChop (https://chopchop.cbu.uib.no), produced by IDT and dissolved in IDTE buffer to a final concentration of 100 µM each. Sequences of the crRNAs were: nphs1_exon_5: GCTCTGACCGTCACATTACG, nphs1_exon_6: CCGACCCACTTGGCACTCGT, nphs1_exon_10: GTGGTATGCATGTCGTACGG, magi2a_AA_exon_1: TAGATCCCCACTCGGACCCC, magi2a_AC_exon_2: CGTGGTCCAAAGAGCCCTTC, magi2a_AB_exon_4: TTACGGCACACCAAAGCCCC. A 5 µM solution of guide RNA was prepared by diluting equal amounts of crRNA and tracrRNA in nuclease-free duplex buffer (IDT) and heating the solution to 95°C for 5 min and stepwise cooling down to 20°C over 10 min in a Mastercycler (Eppendorff). In order to produce RNPs, a 0.5 µg/µl solution of Alt-R® S.p. HiFi Cas9 Nuclease V3 was prepared in sterile PBS, mixed 1:1 with the aforementioned gRNA solution, and incubated at 37°C for 10 min, and stored at 4°C. 1-2 nl of RNP solution was injected in 1-cell stage Tg(fabp10a:gc-eGFP), mitfa^-/-^ roy^-/-^ embryos using an Eppendorff Transjector Turbo, and Femtotips II glass capillaries (Eppendorff). Injected Embryos and respective control groups (negative control targeting an intronic region of the human HBB gene, and non-injected embryos of the same clutch) were reared in 0.5x E3 medium at 28.5°C in the dark. Viability was checked twice daily, and the medium changed once a day. To use only successfully injected embryos for downstream analysis, Atto550 fluorescence was checked at 24 hpf. Embryos with inhomogeneous or negative fluorescence signals were discarded from further analysis. eGFP vessel fluorescence was checked on 4 and 5 dpf using the Acquifer Imaging machine. Larvae were fixed or lysed at 5 dpf for further analysis. PCR amplification and gel electrophoresis of the genomic regions targeted by the RNPs, showed the appearance of additional-sized bands, indicating the presence of insertions or deletions (indels) at the respective binding site. Restriction fragment polymorphism analysis using restriction digestion of the Protospacer Adjacent Motif (PAM) site of the respective RNPs showed either complete failure or decrease of digestion of the PCR products indicating destruction of the PAM due to the presence of indels. PCR products were Sanger-sequenced and analyzed using the inference of CRISPR-edits (ICE) tool that predicts indel frequency in sequencing data which showed >80% knockout efficiency.

### Quantification of vascular fluorescence *in vivo*

The vascular fluorescence intensity measurement of gc-eGFP-expressing larvae was described previously in detail [13]. In brief, larvae were anesthetized with 0.4 mg/ml tricaine (Sigma-Aldrich) and transferred to a 96-well plate prepared with agarose molds. After lateral orientation, overview images were acquired with a 2x objective (2048×2048, 3.25 µm/px) in the Acquifer Imaging Machine (Luxendo GmbH, Heidelberg, Germany). The tail vasculature was focused with a 10x objective (2048×2048, 0.65 µm/px), and a short brightfield movie (10 frames) followed by 1 frame in the 488 channel was acquired. A custom writted FIJI script automatically detects and segments the vasculature based on the movement of erythrocytes in the brightfield movie and measures the eGFP fluorescence intensity exclusively in the vasculature. The output is the mean vascular eGFP fluorescence in the segmented area for each larva in arbitrary greyscale units (16 bit) per px. The mean vascular fluorescence of all larvae from the uninjected group was used to normalize the data and the fluorescence ratios were used as statistical input.

### Zebrafish histology

5 dpf larvae were fixed and sectioned as described before in detail[19]. Briefly, larvae were fixed in 2% paraformaldehyde pH 7 for 1h at RT, immersed in 30% sucrose in PBS for at least 1h at RT, and snap-frozen in a 1:1 mix of OCT (TissueTek, Sakura) and 30% sucrose in PBS. 6 µm transversal sections were obtained on a Leica Leica SM 200R cryomicrotome, collected on SuperFrost microscope slides (VWR) and air-dried at room temperature. Sections were rehydrated in 1x PBS, and permeabilized with 0.1% Triton X-100 (Sigma) in 1x PBS for 30 seconds. Antigenic epitopes were demasked with 1% SDS. Sections were blocked with above mentioned blocking solution. The sections were incubated with primary antibodies (1:1000 in blocking solution, polyclonal rabbit anti-MAGI2 IgG, HPA013650, Sigma; 1:2000 in blocking solution rabbit anti-zebrafish Nephrin, Innovagen, Lund, Sweden) at 4 °C overnight. After three washes in 1x PBS and incubation with the Alexa 647 conjugated goat anti-rabbit F(ab) antibody fragment (1:800) the slides were incubated with 0.013mg/ml Hoechst 33342 (Sigma-Aldrich) for 10 minutes, washed in 1xPBS and mounted in Mowiol for microscopy. Transmission electron microscopy was performed as described before[19].

### Statistical analysis

GraphPad prism V9.3.1 (471) (GraphPad Software, CA, USA) was used for all statistical analyses. Gaussian distribution was checked by Kolmogorov-Smirnov testing. If passed, Student’s t test was used for significance testing. For statistical testing of nonparametric data, Mann-Whitney U test was applied. P-values lower 0.05 were considered statistically significant.

## Acknowledgements

FS was supported by a scholarship of the Gerhard-Domagk Masterclass of the University Medicine Greifswald, Germany. AI was funded by an ERA-EDTA long term fellowship LTF RLTF 1593/2018. The work of the STOP-FSGS consortium was funded by the German Ministry for Science and Education (BMBF, STOP-FSGS-01GM2202A) to NE, PTB, and TBH). TBH was supported by the DFG (CRC1192) and by the European Research Council-ERC (CureFSGS). This work was supported by a grant of the clinical research unit (KFO 329, BR 2955/8-1 to PTB, BE 2212/23-1 and 2212/24-1 to TB).

