## Supplementary figures and images for "The Regulation of MAGI2 and its Diagnostic Application in Podocytopathies"

### Suppl. Fig.

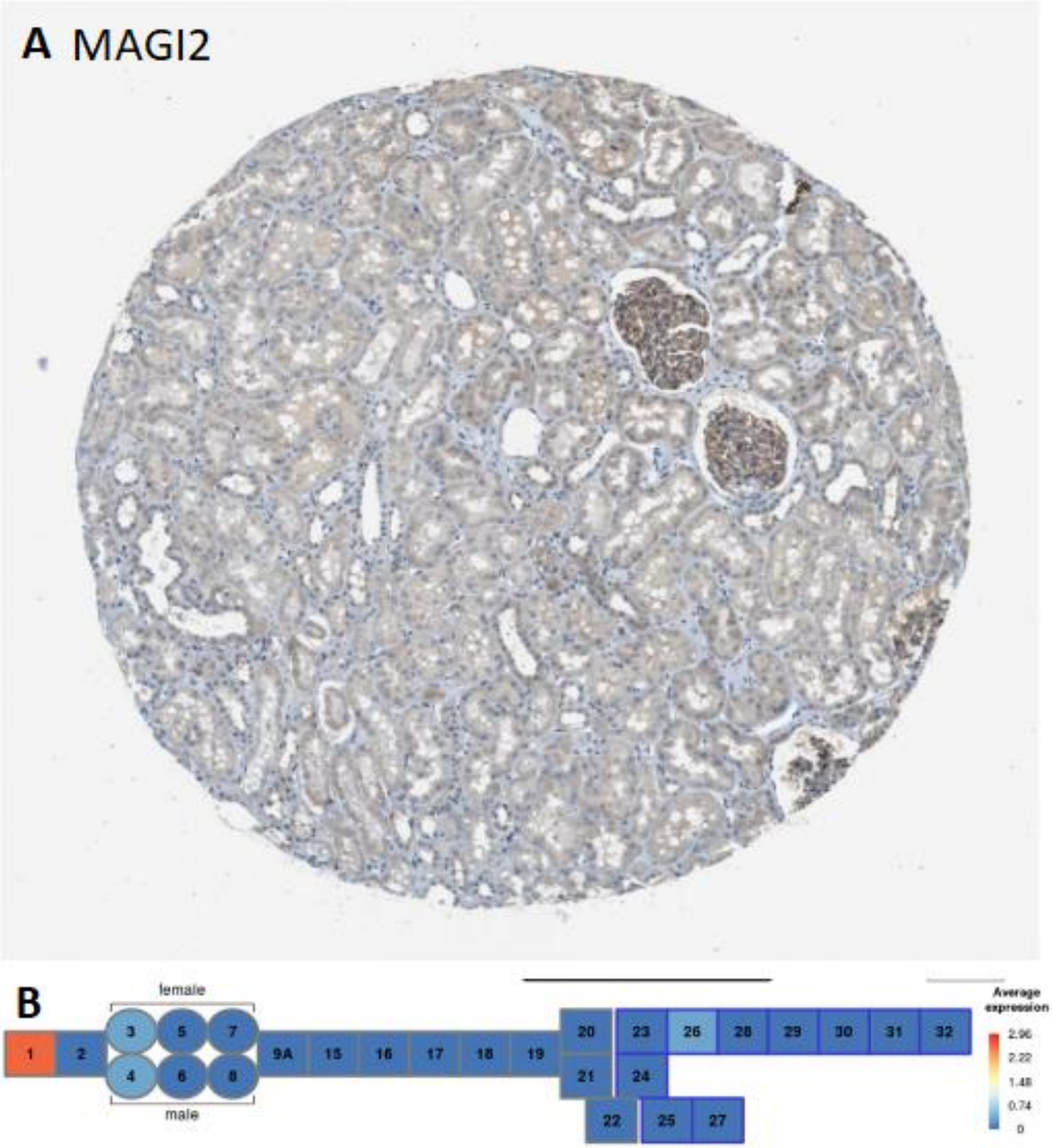

Suppl. Fig. 2

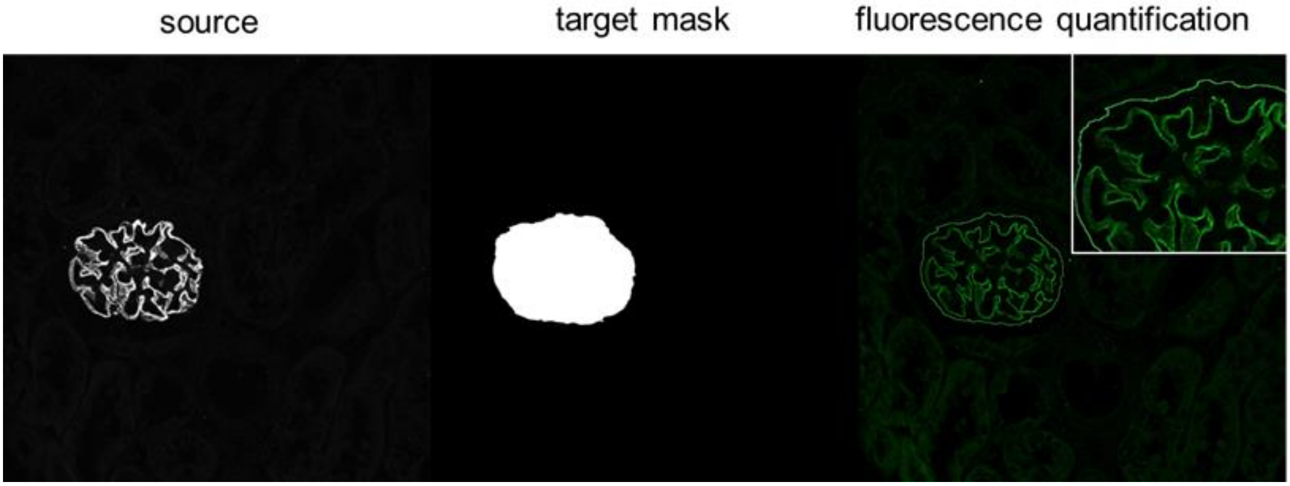
